# White matter microstructure predicts focal and broad functional brain dedifferentiation in normal aging

**DOI:** 10.1101/779264

**Authors:** Jenny R. Rieck, Karen M. Rodrigue, Denise C. Park, Kristen M. Kennedy

## Abstract

Ventral visual cortex exhibits highly organized and selective patterns of functional activity associated with visual processing. However, this specialization decreases in normal aging, with functional responses to different visual stimuli becoming more similar with age, a phenomenon termed “dedifferentiation”. The current study tested the hypothesis that age-related degradation of the inferior longitudinal fasciculus (ILF), a white matter pathway involved in visual perception, could account for dedifferentiation of both localized and distributed brain activity in ventral visual cortex. Participants included 281 adults, ages 20-89, from the Dallas Lifespan Brain Study who underwent diffusion-weighted imaging to measure white matter diffusivity, as well as functional magnetic resonance imaging to measure functional selectivity to viewing photographs from different categories (e.g., faces, houses). In general, decreased ILF anisotropy significantly predicted both focal and broad functional dedifferentiation. Specifically, there was a localized effect of structure on function, such that decreased anisotropy in a smaller mid-fusiform region of ILF predicted less selective (i.e., more dedifferentiated) response to viewing faces in a proximal face-responsive region of fusiform. On the other hand, the whole ILF predicted less selective response across broader ventral visual cortex for viewing animate (e.g., human faces, animals) versus inanimate (e.g., houses, chairs) images. This structure-function relationship became weaker with age and was no longer significant after age 70. These findings indicate that decreased white matter anisotropy is associated with maladaptive differences in proximal brain function and is an important variable to consider when interpreting age differences in functional selectivity.

## Introduction

Ventral visual cortex encompasses regions of occipito-temporal cortex that are uniquely specialized for the identification and recognition of visual stimuli (Grill-Spector & Malach, 2004). Abundant early neuroimaging work has demonstrated selective and localized brain response when viewing certain categories of images (e.g., face-selective regions of fusiform gyrus; Kanwisher, McDermott, & Chun, 1997; Epstein, Harris, Stanley, & Kanwisher, 1999; Haxby et al., 1994), with further evidence for category-specific patterns of activity distributed broadly across ventral visual cortex (Haxby et al., 2001; O’Toole, Jiang, Abdi, & Haxby, 2005). Moreover, one of the largest distinctions in ventral visual cortex is characterized by activity in lateral and superior regions when viewing animate images (e.g., faces, animals, bodies) versus medial and inferior regions when viewing inanimate images (e.g., tools, houses; Grill-Spector & Weiner, 2014; Haxby et al., 2011; Proklova, Kaiser, & Peelen, 2016; Sha et al., 2014). Researchers have proposed that categorical representations in ventral visual cortex are multifaceted and exist at both a focal and broad level with smaller, specialized regions, like face-selective fusiform associated with narrow category distinctions (i.e., “face” vs. “house”) that are nested within larger functional regions involved in superordinate distinctions (i.e., “animate” vs. “inanimate”; Grill-Spector & Weiner, 2014).

With advanced age, these highly specialized and unique representations associated with processing different visual stimuli become less distinct or “dedifferentiated” (Park et al., 2004; Grady et al., 1994; see Koen & Rugg, 2019 for a review). Specifically, with increasing age, localized regions of fusiform specialized for face processing show a broadened response by activating to non-face stimuli (Park et al., 2004; Park et al., 2012). Furthermore, the distributed patterns of brain activity when viewing faces versus other inanimate stimulus categories (e.g., houses or chairs) become more similar in older age (Burianová, Lee, Grady, & Moscovitch, 2013; Carp, Park, Polk, & Park, 2011). Importantly, this neural dedifferentiation has been associated with poorer performance on measures of fluid processing (Park, Carp, Hebrank, Park, & Polk, 2010; Rieck, Rodrigue, Kennedy, Devous, & Park, 2015) and face-matching (Burianová et al., 2013), suggesting that individual differences in brain activity underlying visual perception may account for cognitive aging processes (Park & Reuter-Lorenz, 2009).

One possible explanation for dedifferentiation of functional activity in aging is degradation of the underlying white matter structure. Aging is characterized by dysmyelination and alteration of white matter fiber organization, which, in turn, likely interferes with the transfer and function of neural signals in gray matter (Barnes & McNaughton, 1980; Daselaar et al., 2015). Prior work has examined how measures of white matter microstructure explain age differences in: task-evoked functional response (Burzynska et al., 2013; de Chastelaine, Wang, Minton, Muftuler, & Rugg, 2011; Hakun, Zhu, Brown, Johnson, & Gold, 2015; Madden et al., 2007; Persson et al., 2006; Zhu, Johnson, Kim, & Gold, 2015; see Bennett & Rypma, 2013 and Warbrick, Rosenberg, & Shah, 2017 for reviews), modulation of functional response amplitude (Brown, Hakun, Zhu, Johnson, & Gold, 2015; Webb, Hoagey, Foster, Rodrigue, & Kennedy, 2019), and functional connectivity between cortical regions (Chen, Chou, Song, & Madden, 2009; Davis, Kragel, Madden, & Cabeza, 2012). However, the impact of age-related white matter degradation on highly specialized functional activity (as found in the ventral visual cortex) is unclear.

Here we examine the impact of aging on the relationship between category-selective activations in ventral visual cortex and underlying microstructure of the inferior longitudinal fasciculus (ILF), the principal white matter pathway that connects occipital cortex to inferior and medial temporal brain regions (Catani, Jones, Donato, & Ffytche, 2003). Behavioral studies find that ventral-temporal white matter (including the ILF) is important for a variety of different visually-related cognitive processes including: reading ability (Ellmore et al., 2010; Yeatman, Dougherty, Myall, Wandell, & Feldman, 2012), recognition of visual stimuli (Davis et al., 2009; Sasson, Doniger, Pasternak, & Assaf, 2010; Tavor et al., 2014), and face discrimination (Thomas et al., 2008). Furthermore, previous work has identified ILF fibers that serve as direct connections between face-selective functional regions of ventral visual cortex (Gschwind, Pourtois, Schwartz, Van De Ville, & Vuilleumier, 2012; Saygin, et al., 2012; Pyles, Verstynen, Schneider, & Tarr, 2013; Weiner et al., 2016; Liu, Hildebrandt, Meyer, Sommer, & Zhou, 2020), suggesting that this white matter structure supports the specialized functional activity in ventral visual cortex.

We hypothesized that age-related degradation of ILF would account for functional dedifferentiation during visual processing at both the focal and distributed level. To test this hypothesis we utilized two brain activation measures of dedifferentiation, also referred to as “selectivity”: (1) selectivity for faces in localized face-selective regions of fusiform gyrus (i.e., fusiform face area); and (2) broad selectivity for animacy across the entire ventral visual cortex. In general, functional selectivity describes the degree to which a specific region of cortex shows increased and selective response during one experimental condition (i.e., viewing faces) but not another (i.e., viewing houses). This hypothesis was tested in the Dallas Lifespan Brain Study, with adults ranging in age from 20-89 years, allowing the exploration of the relationship between white matter and neural dedifferentiation across the adult lifespan, including often excluded groups of middle-aged adults and very old age adults (i.e., > 80 years). We also considered that younger adults would vary in ILF integrity and that an effect of decreased anisotropy in ILF would be maintained, even after age was controlled.

## Methods

### Participants

Participants consisted of a subsample from the Dallas Lifespan Brain Study (DLBS) who underwent passive-viewing of categories of photographs during functional magnetic resonance imaging (fMRI) and also had diffusion tensor imaging data collected (DTI; *N* = 299). Participants’ informed consent was obtained in accordance with protocol approved by the University of Texas at Dallas and the University of Texas Southwestern Medical Center. Participants were screened to be right-handed and fluent English speakers with normal or corrected-to-normal vision (at least 20/30), and if necessary, vision was corrected using MRI-compatible corrective lenses during the fMRI session. Participants were additionally screened to be cognitively intact (Mini Mental Status Exam ≥ 26; Folstein, Folstein, & McHugh, 1975) with no history of neurological or psychiatric conditions, head trauma, drug or alcohol problems, or significant cardiovascular disease. Participants with outlier data points (detailed in *Statistical Analysis* section; *N*=18) for each neural measure were identified and excluded from analyses, resulting in a final sample size of 281 (see **Table 1** for sample demographics).

**Table 1.** Sample demographics by decade.

| **Age Range** | **N**<br>**(% female)** | **Age** | **Years of**<br>**Education** | **MMSE** |
| --- | --- | --- | --- | --- |
| 20 – 29 | 41 (65.9%) | 24.37 (2.78) | 15.95 (2.32) | 28.90 (1.26) |
| 30 – 39 | 40 (60.0%) | 34.06 (2.83) | 16.85 (2.35) | 28.63 (1.17) |
| 40 – 49 | 41 (65.9%) | 45.41 (3.19) | 15.82 (1.99) | 28.54 (1.19) |
| 50 – 59 | 37 (73.0%) | 54.63 (2.87) | 16.89 (2.20) | 28.76 (.925) |
| 60 – 69 | 43 (58.1%) | 64.82 (2.88) | 16.88 (2.16) | 28.21 (1.25) |
| 70 – 79 | 39 (66.7%) | 73.34 (2.75) | 16.23 (2.40) | 28.00 (1.29) |
| 80 – 89 | 40 (60.0%) | 83.65 (2.66) | 16.28 (2.35) | 27.30 (1.11) |
| 20 – 89<br>(whole sample) | 281 (64.1%) | 54.23 (19.99) | 16.41 (2.27) | 28.33 (1.27) |

*Note.* Data were analyzed with age as a continuous variable in all analyses. Values represent in the table represent Mean (standard deviation); MMSE=Mini Mental State Exam.

### MRI acquisition

Participants were scanned on a single Philips Achieva 3T whole body scanner equipped with an 8-channel head coil. DTI volumes were acquired with the following parameters: TR = 4410 ms; TE = 51 ms; flip angle = 90 ms; FOV = 224 x 149 x 224; XY matrix: 112 x 110; 50 slices per volume; 30 directions were acquired at b=1000 s/mm^2^, plus a b0 non-diffusion weighted image. Blood-oxygen-level dependent (BOLD) fMRI data were acquired using a T2*-weighted echo-planar imaging sequence with 43 interleaved axial slices (in a 64 × 64 matrix) acquired parallel to the anterior commissure-posterior commissure line with the following parameters: 3.4 × 3.4 × 3.5 mm^3^ voxels, FOV = 220 mm, TE = 25 ms, TR = 2 sec, FA = 80°. High-resolution anatomical images (used to coregister anatomical masks to native diffusion and functional space) were collected with a T1-weighted MP-RAGE sequence with the following parameters: 160 sagittal slices, 1 × 1 × 1 mm^3^ voxels; 256 × 256 × 160 matrix, FOV = 220 mm, TE = 3.76 ms, TR = 8.19 ms, FA = 12°.

### Diffusion tensor imaging

#### DTI processing

Diffusion volumes were processed using the FMRIB’s Diffusion Toolbox (FDT) in FSL (Behrens, Berg, Jbabdi, Rushworth, & Woolrich, 2007; Behrens et al., 2003). First, volumes underwent eddy current correction to correct for distortions caused by eddy currents in the gradient coils, as well as simple head motion during image acquisition by using an affine registration to a reference volume (using FLIRT). Second, the images were skull stripped using Brain Extraction Tool. Next, diffusion tensors were fit on corrected data using *dtifit* to create a voxel-wise map of the three primary diffusion directions (i.e., λ_1_, λ_2_, and λ_3_) for each participant.

The three diffusion directions were used to compute our primary measure of white matter microstructure: fractional anisotropy (FA), which describes the ratio of diffusion in the primary direction (λ1 or axial diffusivity; AD) relative to perpendicular directions (λ_2_ and λ_3_ or radial diffusivity; RD). Because FA is a composite index of diffusivity in all directions, it is important to consider axial and radial diffusivity interpreting differences in white matter microstructure (e.g., decreased axial diffusivity coupled with increased radial diffusivity could result in decreased FA). Therefore, in the current study significant effects of FA were further explored by also examining the contribution of AD and RD to the statistical models.

#### Isolating white matter structures

The primary white matter pathway of interest was the inferior longitudinal fasciculus (ILF), and we also isolated a smaller mid-fusiform portion of ILF for follow-up analyses. Four additional white matter structures were identified to serve as control white matter regions: the superior longitudinal fasciculus (SLF), the uncinate fasciculus (UF), and the genu and splenium of the corpus callosum.

Probabilistic tractography was conducted to isolate the three bilateral association white matter pathways (ILF, SLF and UF), separately in each participant’s native space using FDT in FSL (Behrens et al., 2007). First, diffusion parameters were estimated at each voxel using *bedpostX*, which generates a probability distribution function of the primary diffusion directions. Then, to determine connectivity between voxels, *probtrackX* was used to estimate the distribution of connections between the seed regions (i.e., origin) and waypoint (i.e., inclusion) regions (described below). The output from this step was a connectivity value for each voxel that represents the number of fibers that passed through that voxel.

All masks were constructed in standard Montreal Neurological Institute (MNI) space and then backwarped to each participant’s native diffusion space; and all seed and waypoint regions were rectangular masks: 6 mm deep (anterior-posterior/coronal); 18 mm wide (left-right/sagittal) and 28 mm high (dorsal-ventral/axial), constructed separately for the left and right hemispheres. A midline exclusion mask was also included to eliminate fibers that crossed hemisphere.

ILF seed masks (i.e., tract origins) were created in occipital white matter, 24 mm anterior to the most posterior slice of the brain, and a single waypoint mask was created in temporal white matter, 50 mm anterior to the seed mask. Seed and waypoint masks were positioned such that they were centered around anterior-posterior oriented white matter as evident on the MNI template (see **Figure 1A** for an illustration of ILF masks and the resulting pathway for a representative participant in cyan). For the SLF, seed masks were constructed in superior white matter on the most posterior coronal slices in which the splenium of the corpus callosum was visible on the MNI template. Two SLF waypoint masks were constructed 22 mm and 44 mm anterior to the seed mask. For the UF, seed masks were positioned in anterior temporal white matter, 30 mm posterior to the tip of the temporal pole. One waypoint mask was placed in the white matter of the anterior floor of the external capsule.

**Figure 1.**
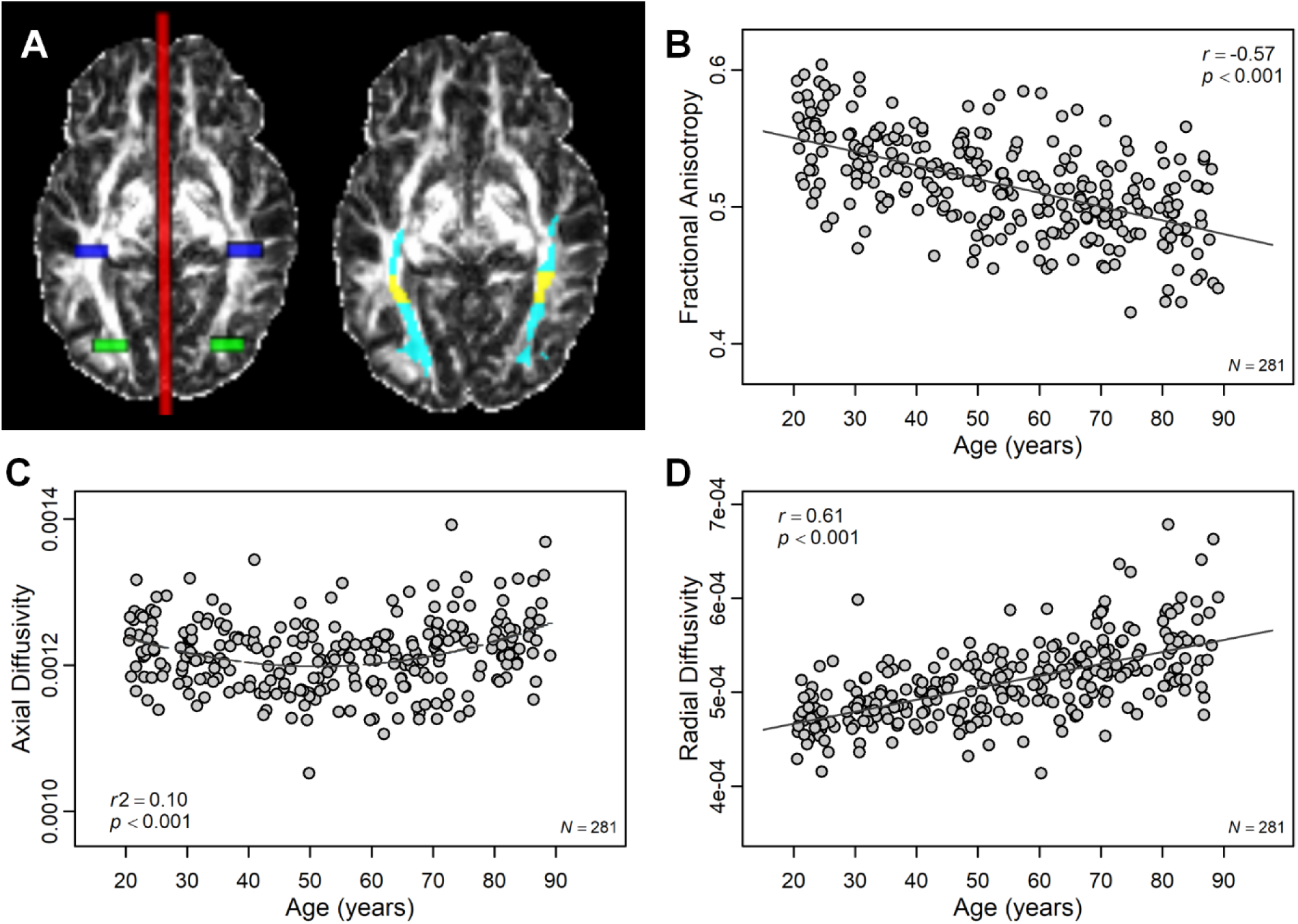
Effect of age on white matter diffusion metrics in ILF. (A) Seed (green), waypoint (blue), and exclusion (red) masks used for probabilistic tractography of left and right ILF (left side) as well as the resulting ILF tract (cyan; right side) are illustrated for a representative participant. Mid-fusiform ILF has been highlighted in yellow. (B) With increasing age, diffusivity in ILF was more isotropic (i.e., lower fractional anisotropy) which was characterized by (C) non-linear age-differences in axial diffusivity and (D) linear age increases in radial diffusivity.

The resulting white matter tracts were thresholded at 15% of the maximum connectivity value, which ensured that tracts only included those voxels with a high likelihood of being connected to the seed and waypoint regions (following Bennett, Motes, Rao, & Rypma, 2012). All tracts were then visually inspected to verify that the backwarping of the seed and waypoint regions to native diffusion space resulted in proper tracking. Thresholded tracts were binarized, and mean FA, AD, and RD were extracted and averaged across hemispheres.

To localize fibers within the genu (anterior portion) and splenium (posterior portion) of the corpus callosum, rather than including pericallosal fibers, regions of interest were hand-traced on the T2-weighted (b0) baseline image for each participant in their native diffusion space with high reliability (intraclass correlation coefficient > .95). Each region was traced with a stylus on an LCD digitizing tablet using Analyze v12.0 software (Mayo Clinic, Rochester, MN). The genu and splenium were both traced on the slices in which they were optimally visible, for each participant with at least three slices included per structure. For each participant, diffusivity measure (e.g., FA, AD, RD) were extracted from the hand-traced genu and splenium and averaged across slices for each structure.

Finally, a smaller portion of ILF in mid-fusiform (proximal to the functional fusiform face area) was identified in order to test regional specificity of structure-function relationships. Because the decision to examine mid-fusiform ILF was *post hoc*, the region was identified in an unbiased manner via a group-level analysis of the fMRI data. Specifically, rectangular masks were created in MNI space (28mm ×12 mm × 18 mm, similar to the ILF tractography seed and waypoint masks) that were centered around the left (Y = -43) and right (Y = -48) Y values for the peak clusters from a group-level analysis of functional activity for faces > houses (illustrated in **Figure 2C**). These MNI masks were then backwarped to each participant’s native diffusion space and overlaid with their ILF pathway generated from tractography. Mean FA was extracted from the conjunction of the backwarped mask and the ILF pathway in order to create a measure of mid-fusiform ILF microstructure in roughly equivalent anatomical regions for each participant (confirmed via visual inspection) that was proximal to functional activity in fusiform face area. A visualization of this mid-fusiform segment for one participant can be found in **Figure 1A** (yellow portion of ILF).

**Figure 2:**
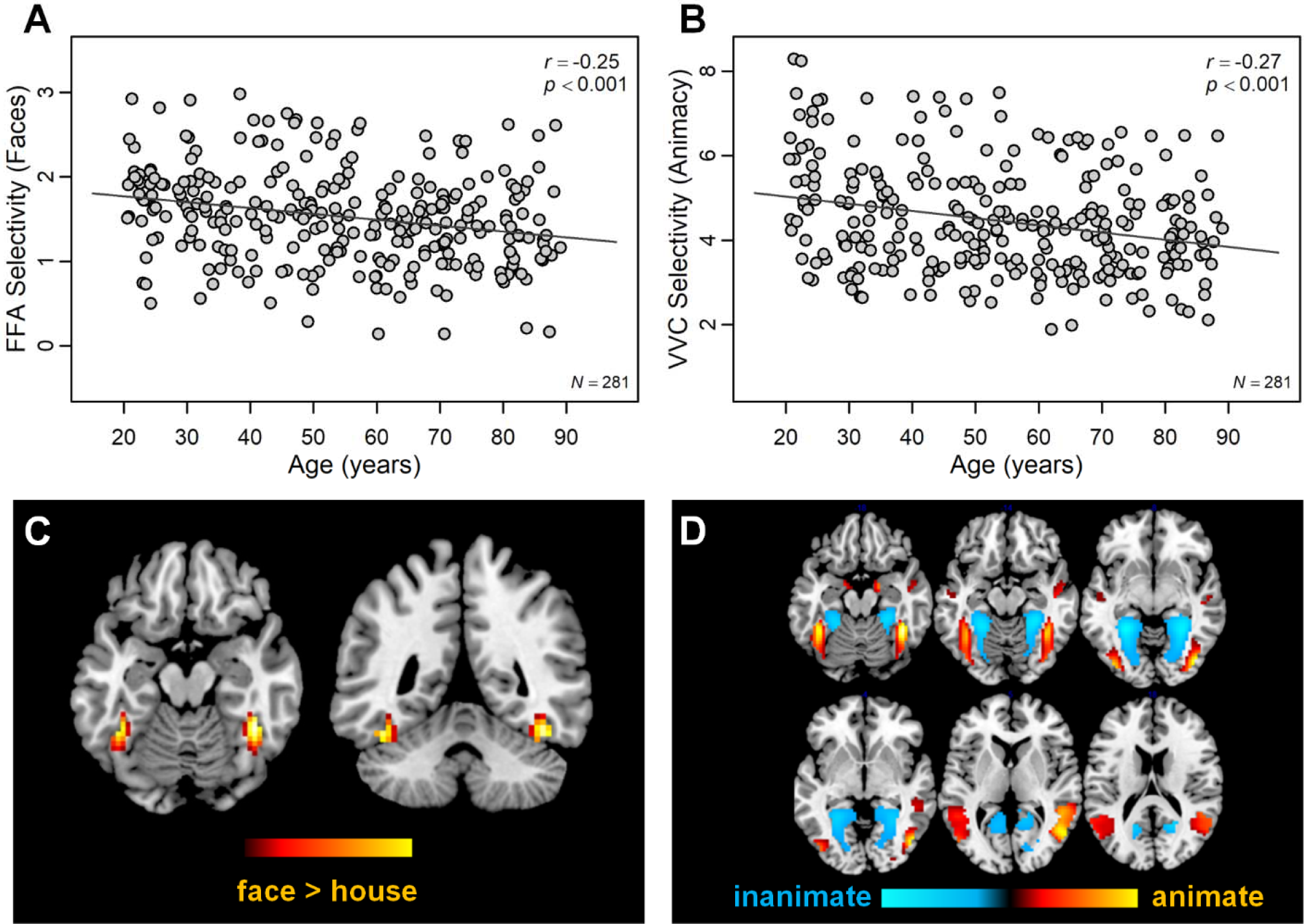
Effect of age on functional selectivity. (A) With increasing age, selectivity for faces in fusiform face area decreased. (B) Likewise, selectivity for animacy across ventral visual cortex (VVC) decreased. Functional activation maps illustrate (C) fusiform face area and (D) activity associated with viewing animate versus inanimate images mapped to a common template space.

### Functional magnetic resonance imaging

#### Visual stimuli and task design

While in-scanner, participants were instructed to view images from six categories: human faces, primates, cats, houses, chairs, and phase-scrambled control stimuli. Each category was composed of 64 grayscale photographs (400 pixels wide x 300 pixels tall), and cat stimuli were made up of 32 domestic cat and 32 wild cat photographs. The images of human faces were taken from our published face library (Kennedy, Hope, & Raz, 2009; Minear & Park, 2004). Animal photographs (primates, domestic cats, and wild cats) were taken from the internet and cropped so that the animal’s face was clearly visible. Like the human faces, only animal images with front-facing and neutral expressions were chosen. Houses were photographed from various locations across the United States. The photographs of chairs were taken from furniture websites. Control images were created by scrambling the phase information in all the experimental stimuli so that the spatial frequency information was preserved, but the visual information was meaningless.

Visual stimuli were presented in a block design across two seven-minute runs. There was a 10-sec fixation screen at the beginning and end of each run to allow brain tissue magnetization to reach a steady state and to allow for participants to become acclimated to the noise of the scanner. Within each run, 24 blocks (four blocks from the six categories) were presented in pseudorandom order. Within each block there were eight images randomly selected from the same category so that each image was only presented one time. Individual images were presented for 2 seconds each with no inter-stimulus interval. Participants were not required to make any response during the scan, but were instructed to pay attention to each image. A subset of data from this task (i.e., the human and house viewing conditions) have been discussed in previous work (Park et al., 2012).

#### fMRI processing

Individual participant’s time series data were preprocessed with Statistical Parametric Mapping 8 (SPM8; Wellcome Department of Cognitive Neurology, London, UK). First, images were corrected for differences in slice time acquisition. Next, individual volumes were corrected for within-run participant movement. Finally, images were smoothed with an isotropic 8 mm^3^ full-width-half-maximum Gaussian kernel. For the current study, the primary analysis of the fMRI data was conducted in participant native-space to ensure the least amount of processing prior to analysis. However, an additional pipeline was run which included a spatial normalization to a standardized MNI template prior to the smoothing step in order to ensure that warping the data to a standard space did not differ from the analysis conducted in native space.

First-level statistics were conducted using the general linear model in SPM8 to model BOLD response to each of the six image categories viewed in scanner: human faces, primates, cats, houses, chairs, and scrambled. For the current study, additional first-level models were conducted to estimate neural response to animate (human, primate, and cat) and inanimate (house and chair) categories. In all models, motion parameters for each individual were also included as nuisance regressors to control for individual differences in movement across the scanning session. The resulting parameter estimates (β-weights) from the first-level models quantified how well functional activity in each voxel corresponded to a predicted neural response for that condition, thereby providing an estimate of BOLD response to each experimental condition.

### Quantifying functional measures of dedifferentiation

For the current study, two indices of functional dedifferentiation, also referred to as “selectivity”, were examined: (1) functional selectivity for faces in face-selective regions of fusiform gyrus (e.g., fusiform face area); and (2) broad functional selectivity for animacy across the entire ventral visual cortex.

To quantify face selectivity, we focused on activation patterns in the fusiform face area (FFA), as this is the region of the core face-network that shows the strongest age-related dedifferentiation compared to other regions of the face network (e.g., occipital face area, superior temporal sulcus; Park et al., 2012). Following Park and colleagues (2012), for each participant, left and right FFA were identified within participant-native space by isolating a ∼200 mm^2^ contiguous region around the peak voxel in fusiform gyrus that evoked significant response to viewing human faces versus scrambled images (FFA voxels that have been mapped to a common space are illustrated in **Figure 2C**). Within the functionally defined FFA, neural selectivity was quantified as the average difference in neural response to human faces compared to houses (more details can be found in Park et al., 2012). For the current study, FFA selectivity measurements were averaged for left and right hemispheres.

A second index of selectivity was also computed based on prior work demonstrating widespread differences in functional representations associated with animacy (Grill-Spector & Weiner, 2014; Haxby et al., 2011; Proklova et al., 2016; Sha et al., 2014). For the current study, selectivity for animacy was quantified by calculating the similarity between neural response to animate images (human, primate, and cat) and inanimate images (houses and chairs) using Euclidian distance, which is a measure of the magnitude of difference in neural response to different conditions across many voxels (Kriegeskorte, Mur, & Bandettini, 2008); a group-level contrast of animacy mapped to a common space is illustrated in **Figure 2D**).

Ventral visual cortex was identified for each participant using a participant-specific anatomical mask in conjunction with a functionally derived mask that isolated voxels in the ventral visual mask that showed significant response to any of the visual stimuli. Specifically, an anatomical mask of the ventral visual pathway was backwarped to participant native-space using bilateral regions derived from the Automated Anatomical Labeling template (AAL; Tzourio-Mazoyer et al., 2002), specifically: calcarine sulcus, cuneus, lingual gyrus, inferior occipital gyrus, fusiform gyrus, parahippocampal gyrus, inferior and middle temporal gyrus, and temporal. Although not all of these regions are necessarily part of the “ventral visual pathway”, they were selected to encompass the large swathe of cortical regions to which ILF projects in addition to functional regions involved in visual processing. Second, a functional mask was generated for each individual to isolate voxels within the ventral visual pathway that were responsive to visual input. Following O’Toole and colleagues (2014), the functional mask was defined as those voxels within the ventral visual cortex that varied significantly across the six stimulus conditions as determined by an analysis of variance *F*-test. For the current study, the significance threshold was set to *p* < .00001 (O’Toole et al., 2014). Relaxing or increasing the significance threshold for the functional masks did not alter the results reported here.

### Statistical analysis

Outliers were removed from the sample by identifying those participants who evidenced outlier data points for any of the three primary brain variables of interest: ILF FA; FFA face selectivity; and ventral visual animacy selectivity. To ensure outlier removal was not influenced by normal aging effects, first age was regressed out of each brain variable of interest. Outliers were identified as those data points outside 1.5 times the interquartile range for the standardized residuals (after accounting for age). Eighteen total outliers were identified (ILF FA, *n* = 5; face selectivity, *n* = 7; animacy selectivity outliers, *n* = 5; both face and animacy selectivity, *n* = 1), for a final sample of *N* = 281.

Using R (R Core Team), several hierarchical regression models were conducted in order to examine the predictive effect of white matter microstructure on the different functional measures of selectivity. For each hierarchical regression, age was entered first, followed by FA for the control white matter regions (i.e., SLF, UF, genu, and splenium), and finally, ILF FA (the white matter pathway of interest). This analysis allowed us to determine if ILF microstructure predicted differences in functional selectivity beyond the effect of aging and controlling for white matter FA in other brain regions. Because the animacy measure was derived across a large region of ventral visual cortex in each individual’s native space, the size of each individual’s ventral visual area (as computed through their anatomically-derived mask; range = 4769-7744 voxels in size; mean number of voxels = 5893) was also included as a nuisance covariate in each model to ensure that results were not influenced by individual differences in ventral visual size.

The regression models also allowed us to examine the shared (i.e., *R*^*2*^) and unique (i.e., η^2^) variance for each of the predictors to understand the age and ILF microstructure contributions to the overall model. For the highest-level regression model, we also included age × ILF microstructure interactions to examine how age might be modulating the relationship between white matter microstructure and functional selectivity. For models in which mean FA in ILF was found as a significant predictor, additional regressions were conducted (with predictors from the highest-level significant hierarchical regression model) using other indicators of white matter diffusivity (i.e., axial and radial diffusivity) in order to determine which diffusion metrics accounted for the effect of FA.

## Results

### Age Effects on Structural and Functional Measures

The association of each structural and functional measure to age was individually assessed to establish that the usual sensitivity to aging was observed for each marker. As expected, with increasing age, diffusion in ILF was more isotropic as evident by the decrease in FA values (*r* = -.57, *p* < .001), suggesting that integrity in ILF degrades with increasing age (**Figure 1B**). Age-related decreases in FA were characterized by a quadratic effect of age on axial diffusivity (*r*^*2*^ = .10, *p* < .001; **Figure 1C**) and linear effect of age on radial diffusivity (*r* = -.61, *p* < .001; **Figure 1D**). Specifically, participants age 20 to 51 show age-related increases in radial diffusivity (*r* = .31, *p* < .001) and slight decreases in axial diffusivity (*r* = -.32, *p* < .001). On the other hand, participants ages 52 to 89 showed age-related increases in both radial (*r* = .42, *p* < .001) and axial diffusivities (*r* = .30, *p* < .001), which might be suggestive of a more severe pattern of degradation in adults over the age of 52 (Bennett, Madden, Vaidya, Howard, & Howard, 2010; Burzynska et al., 2010).

Both functional measures of selectivity showed decreases with increasing age (**Figure 2**). Specifically, in face-selective regions of fusiform gyrus (i.e., FFA), the difference in neural response to faces compared to houses was decreased in older age (*r* = -.25, p < .001; **Figure 2A**). Likewise, across the ventral visual cortex, the similarity of response between animate and inanimate images was decreased in older age (*r* = -.27, *p* < .001; **Figure 2B**).

### Structure-Function Associations

Hierarchical regression models were conducted in which age and white matter metrics and nuisance covariates were entered to predict the two different measures of functional selectivity (i.e., face selectivity or animacy).

### Face Selectivity

The first set of models predicted selectivity for faces versus houses in FFA and found that older age predicted decreased face selectivity (Δ*R*^*2*^ = .062, *p* < .001). However, adding mean FA of the ILF to the model (after controlling for age and FA in non-preferred white matter pathways) did not significantly increase the amount of variance explained by the model (Δ*R*^*2*^ = .002, *p* = .42; **Table 2A**).

**Table 2.** Regression models: Predicting selectivity for faces in fusiform face area.

| A. Fractional Anisotropy (FA) |  |  |  |  |  |  |  |  |
| --- | --- | --- | --- | --- | --- | --- | --- | --- |
| Model | Steps | Measurement | $R^2$ | $\Delta R^2$ | $\Delta R^2$ $p$-value | $F$-value | $p$-value | $\eta^2$ |
| 1 | 1 | Age | .062 | - | - |  |  |  |
| 2 | 1 | Age | .068 | .006 | .797 |  |  |  |
|  | 2 | Other white matter FA<br>(SLF, UF, Genu, Splenium) |  |  |  |  |  |  |
| 3 | 1 | Age | .070 | .002 | .421 |  |  |  |
|  | 2 | Other white matter FA |  |  |  |  |  |  |
|  | 3 | Whole ILF FA |  |  |  |  |  |  |
| 4 | 1 | Age | .092 | .022 | .012 | 1.73 | .001 | .038 |
|  | 2 | Other white matter FA |  |  |  | all < 1 | all > .1 | all < .003 |
|  | 3 | Whole ILF FA |  |  |  | 2.02 | .156 | .007 |
|  | 4 | Middle Fusiform ILF FA |  |  |  | 6.43 | .012 | .023 |
| B. Radial Diffusivity (RD) |  |  |  |  |  |  |  |  |
| Model | Steps | Measurement | $R^2$ | $\Delta R^2$ | $\Delta R^2$ $p$-value | $F$-value | $p$-value | $\eta^2$ |
| 1 | 1 | Age | .095 | - | - | 14.73 | < .001 | .051 |
|  |  | Other white matter RD |  |  |  | all < 1 | all > .1 | all < .007 |
|  |  | Whole ILF RD |  |  |  | 2.84 | .093 | .010 |
|  |  | Middle Fusiform ILF RD |  |  |  | 3.97 | .047 | .015 |
| C. Axial Diffusivity (AD) |  |  |  |  |  |  |  |  |
| Model | Steps | Measurement | $R^2$ | $\Delta R^2$ | $\Delta R^2$ $p$-value | $F$-value | $p$-value | $\eta^2$ |
| 1 | 1 | Age | .099 | - | - | < 1 | .588 | .001 |
|  |  | Other white matter AD |  |  |  | all < 1 | all > .1 | all < .001 |
|  |  | Whole ILF AD |  |  |  | < 1 | .619 | .015 |
|  |  | Middle Fusiform ILF AD |  |  |  | 15.24 | < .001 | .053 |

*Note.* $F$ -, $p$ - and $\eta^2$ are reported for the highest-step significant regression models; other white matter pathways (SLF, UF, Genu and Splenium) were entered as separate predictors; SLF=Superior Longitudinal Fasciculus; UF=Uncinate Fasciculus; ILF=Inferior Longitudinal Fasciculus.

Given that the ILF is a fairly long fiber pathway, (i.e., up to 10 cm in length; Catani et al., 2003) and the FFA regions identified in the current study were roughly 8 mm spheres, it is possible that the size-scale difference between the two measures accounted for the lack of the relationship between FA and face selectivity in FFA. Therefore, an additional regression model was conducted to determine if FA in a more local (i.e., mid-fusiform) portion of ILF was a better predictor of selectivity than FA averaged across the whole ILF pathway (see Methods for details on how this region was delineated). Specifically, left and right mid-fusiform FA were averaged and entered in a hierarchical regression model (after age, control regions FA, and whole ILF FA) to predict selectivity for faces within FFA.

This analysis revealed that decreased FA within the mid-fusiform region of ILF predicted decreased face selectivity within FFA (β = 3.33, *F*(1,273) = 6.43, *p* = .012, η^2^ = .023), even when accounting for age, other white matter regions, and whole ILF FA (Δ*R*^*2*^ = .022, *p* = .012; see **Figure 3** for an illustration of this association).^1^ Examining the unique variance (i.e., η^2^) explained by each predictor in this model revealed that age accounted for 3.8% and mid-fusiform FA accounted for 2.3% of the total model variance (9.2%). These findings indicate that even though both age and white matter have unique contributions to individual differences in FA selectivity, together these variables explain more variance than alone (**Table 2A**). Including an interaction term between age and mid-fusiform FA for this final model did not result in a significant interaction (*F*(1,272) = .266, *p* = .606), nor did it explain more variance than the previous model (Δ*R*^*2*^ = .001, *p* = .606).

**Figure 3.**
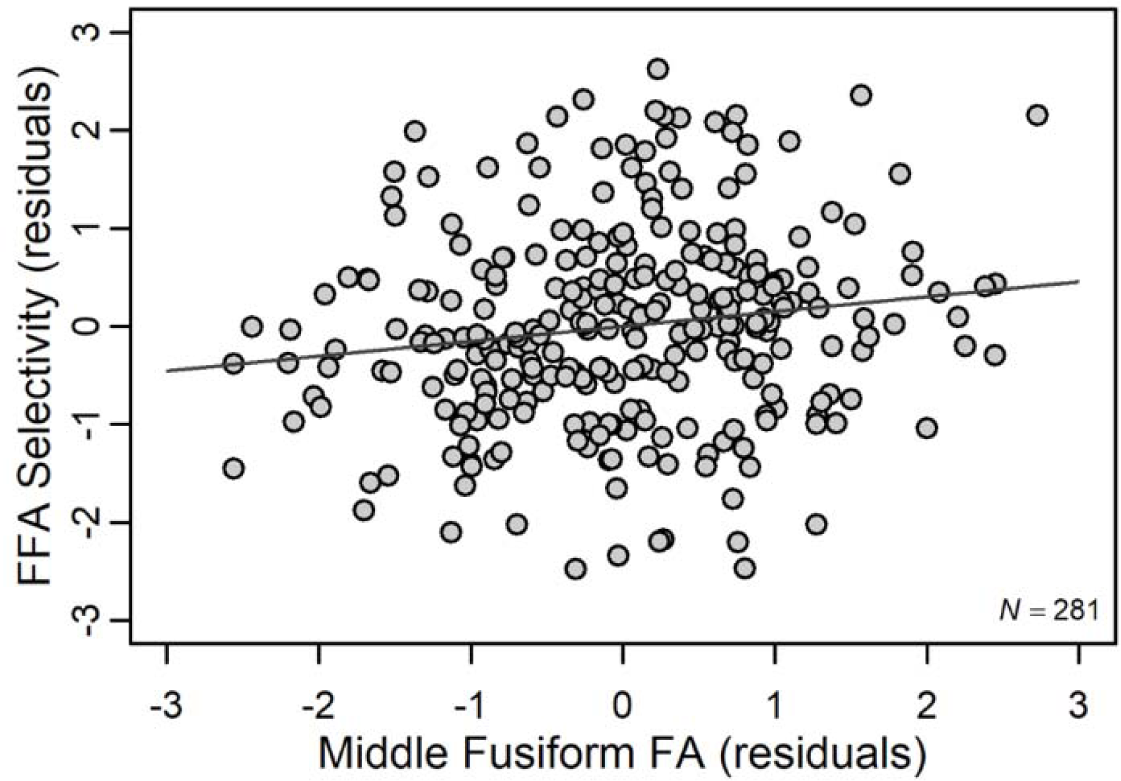
Effect of mid-fusiform FA on FFA Selectivity. Increased FA in a portion of middle-fusiform ILF corresponded to increased face-selectivity in fusiform face area (FFA). Standardized residuals (after accounting for age and other white matter tracts) are plotted for middle-fusiform FA and FFA selectivity

Two additional regression models were conducted using axial and radial diffusivity, respectively, for each white matter region. These models demonstrated that the association between decreased white matter microstructure in mid-fusiform and decreased face selectivity in FFA was characterized by increased radial (*R*^*2*^ =.095, *F*(1,273) = 3.97, *p* = .047, η^2^ = .015; **Table 2B**) and decreased axial diffusivities (*R*^*2*^ =.099, *F*(1,273) = 15.24, *p* < .001, η^2^ = .053; **Table 2C**) when accounting for age and control region diffusivity measures.

### Animacy

The second set of hierarchical regression models predicted selectivity for animacy across ventral visual cortex. As expected, age was a significant predictor of selectivity for animacy (Δ*R*^*2*^ = .071, *p* < .001). Adding FA in control white matter regions (ΔR^2^ = .006, *p* = .788) and ventral visual volume size (Δ*R*^*2*^ = .001, *p* < .583) did not explain significantly more variance in neural selectivity. When accounting for all of these variables, whole ILF FA remained a significant predictor of neural selectivity for animacy (β = 10.70, Δ*R*^*2*^ = .037, *p* < .001; **Table 3A**); specifically, decreased white matter FA in ILF predicted decreased selectivity across ventral visual cortex.

**Table 3.** Regression models: Predicting selectivity for animacy across ventral visual cortex.

| **A. Fractional Anisotropy (FA)** |  |  |  |  |  |  |  |  |
| --- | --- | --- | --- | --- | --- | --- | --- | --- |
| Model | Steps | Measurement | $R^2$ | $\Delta R^2$ | $\Delta R^2 p$-value | $F$-value | $p$-value | $\eta^2$ |
| **1** | 1 | Age | **.071** | - | - |  |  |  |
| **2** | 1 | Age | .077 | .006 | .788 |  |  |  |
|  | 2 | Other white matter FA<br>(SLF, UF, Genu, Splenium) |  |  |  |  |  |  |
| **3** | 1 | Age | .078 | .001 | .583 |  |  |  |
|  | 2 | Other white matter FA |  |  |  |  |  |  |
|  | 3 | Ventral visual cortex size |  |  |  |  |  |  |
| **4** | 1 | Age | **.115** | **.037** | **.001** |  |  |  |
|  | 2 | Other white matter FA |  |  |  |  |  |  |
|  | 3 | Ventral visual cortex size |  |  |  |  |  |  |
|  | 4 | ILF FA |  |  |  |  |  |  |
| **5** | 1 | Age | **.130** | **.015** | **.031** | 3.75 | .054 | .014 |
|  | 2 | Other white matter FA |  |  |  | all < 1 | all > .1 | all < .008 |
|  | 3 | Ventral visual cortex size |  |  |  | < 1 | .545 | .001 |
|  | 4 | ILF FA |  |  |  | **12.44** | **<.001** | **.044** |
| | 5 | Age × ILF FA | | | | **4.66** | **.032** | **.017** |
| **B. Radial Diffusivity (RD)** |  |  |  |  |  |  |  |  |
| Model | Steps | Measurement | $R^2$ | $\Delta R^2$ | $\Delta R^2 p$-value | $F$-value | $p$-value | $\eta^2$ |
| **1** | 1 | Age | **.137** | - | - | **5.90** | **.016** | **.021** |
|  |  | Other white matter RD |  |  |  | all < 1 | all > .1 | all < .004 |
|  |  | Ventral visual cortex size |  |  |  | < 1 | .395 | .003 |
|  |  | ILF RD |  |  |  | **10.52** | **.001** | **.037** |
| | | Age × ILF RD | | | | **4.51** | **.035** | **.016** |
| **C. Axial Diffusivity (AD)** |  |  |  |  |  |  |  |  |
| Model | Steps | Measurement | $R^2$ | $\Delta R^2$ | $\Delta R^2 p$-value | $F$-value | $p$-value | $\eta^2$ |
| **1** | 1 | Age | **.118** | - | - | **25.40** | **<.001** | **.085** |
|  |  | Other white matter AD |  |  |  | all < 1 | all > .1 | all < .003 |
|  |  | Ventral visual cortex size |  |  |  | < 1 | .967 | < .001 |
|  |  | ILF AD |  |  |  | < 1 | .527 | .002 |
| | | Age × ILF AD | | | | < 1 | .762 | < .001 |

*Note.* $F$ -, $p$ - and $\eta^2$ are reported for the highest-step significant regression models; other white matter pathways (SLF, UF, Genu and Splenium) were entered as separate predictors; SLF=Superior Longitudinal Fasciculus; UF=Uncinate Fasciculus; ILF=Inferior Longitudinal Fasciculus.

Adding an age × ILF FA term to this model again explained more variance than the previous step (Δ*R*^*2*^ = .015, *p* = .031), and also resulted in a significant interaction (*F*(1,272)=4.66, *p* = .032; **Table 3A**), indicating that the association between ILF white matter and broad functional selectivity to animacy was age-dependent. Because both age and ILF FA were continuous variables, we used a simple slopes analysis (Preacher et al., 2016; Johnson & Fay, 1950) to examine how age modulated the relationship between structure and function. Specifically, we computed the slope of the relationship between ILF FA and ventral visual selectivity for the mean age in our sample (54 years), as well as 1 standard deviation above (74 years) and below (34 years). The simple slopes analysis revealed that strength of the relationship between whole ILF FA and broad functional selectivity became weaker with increasing age (β_*Age 34*_ = 15.33; β_*Age 54*_ = 10.77; β_*Age 74*_ = 6.21; **Figure 4A**). Furthermore, using the Johnson-Neyman approach to examine the structure-function slopes across the entire lifespan confirmed that the relationship between ILF FA and functional selectivity was no longer significant after about age 70 (**Figure 4B**, dotted vertical line)

**Figure 4.**
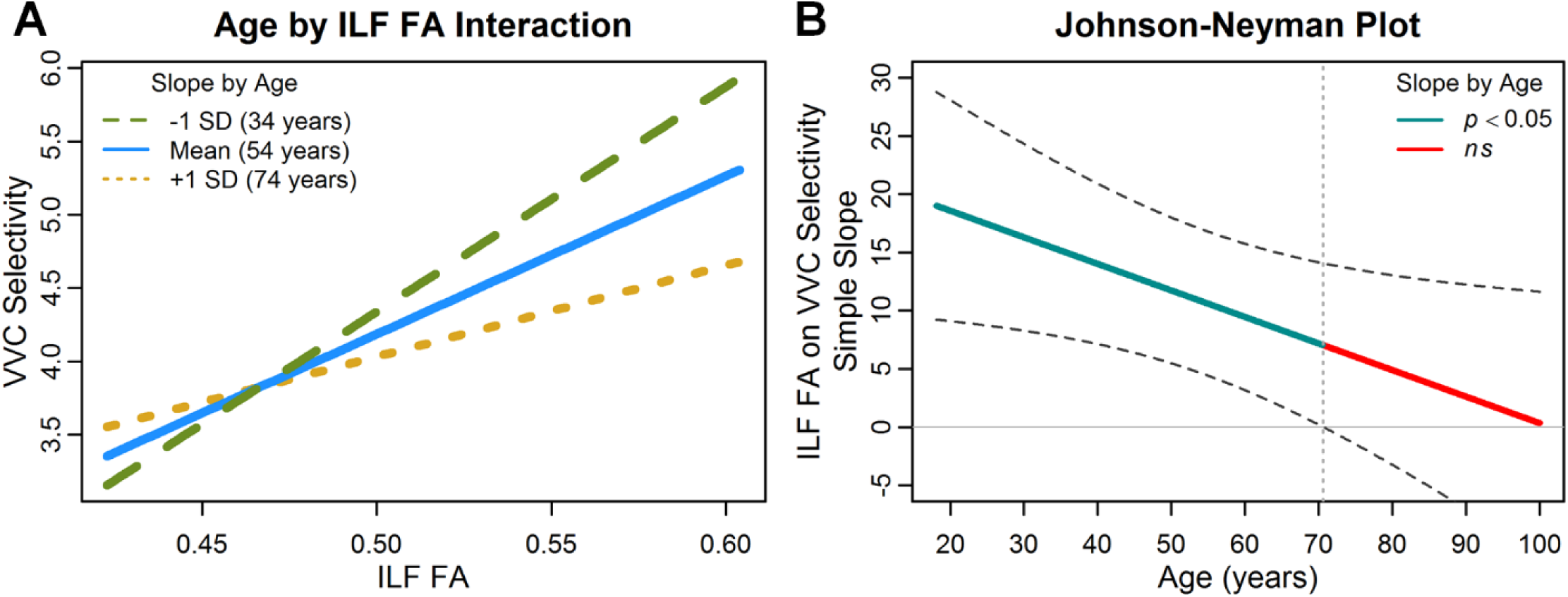
Effect of ILF FA on ventral visual selectivity to animacy depends on age. (A) Increased FA in whole ILF corresponded to increased selectivity for animacy across ventral visual cortex (VVC). However, this relationship weakened with age, as evident by plotting the slope of the relationship between ILF FA and VVC selectivity for the sample mean age (54 years; solid blue line) and 1 standard deviation (SD) above (74 years; dotted yellow line) and 1 SD below (34 years; dashed green line). (B) The Johnson-Neyman plot illustrates the structure-function slopes at each point of the lifespan with 95% confidence intervals (dashed lines) indicating that the ILF FA and VVC selectivity relationship was no longer significant after age 70 (where the confidence intervals cross 0, delineated with a vertical dotted line).

Two additional regression models were conducted using axial and radial diffusivity for each white matter region. These models indicated that decreased selectivity for animacy was predicted by increased radial diffusivity (*R*^*2*^ =.137, *F*(1,272) = 10.52, *p* = .001, η^2^ = .037; **Table 3B**), with no relationship found for axial diffusivity (*R*^*2*^ =.118, *F*(1,272) < 1, *p* = .527, η^2^ = .002; **Table 3C**). As with FA, the age × ILF radial diffusivity interaction was significant (*F*(1, 272)= 4.51, *p* = .035, *η*^*2*^ = .016). A simple slopes analysis of the age × radial diffusivity interaction found that the relationship between white matter microstructure and ventral visual selectivity weakened with age (*β*_*Age 34*_ = -13772; *β*_*Age 54*_ = -9750; age 74: *β*_*Age 74*_ = -5727) and became non-significant after age 72.

## Discussion

This work provides new evidence of a direct relationship between ILF microstructure properties and functional selectivity in ventral visual cortex during visual perception. Specifically, decreased white matter anisotropy in ILF corresponded to less selective activation for animacy across the ventral visual cortex, independent of the effect of age. This effect was not found when trying to predict neural selectivity for faces within a specialized region of the face processing network. However, white matter anisotropy within a smaller and more localized portion of the ILF did predict functional selectivity in fusiform face area. These results suggest that whole ILF white matter may be a stronger predictor of broad differences in selectivity across the entire ventral visual cortex, whereas more focal regions of ILF may be a stronger predictor of dedifferentiation of specialized face processing.

### Whole ILF microstructure predicts broad but not focal measures of neural selectivity

The ILF is a fairly long fiber pathway that interconnects many areas of cortex between the occipital and temporal poles (Catani et al., 2003)—essentially along what is considered the ventral visual stream—and it is clear from behavioral studies that the ILF is involved in variety of visually-dependent cognitive processes including: reading ability (Ellmore et al., 2010; Yeatman et al., 2012), recognition of visual stimuli (Davis et al., 2009; Sasson, et al., 2010; Tavor et al., 2014), speed of processing (Bennett et al., 2012; Turken et al., 2008). Yet, it is only more recently that researchers have examined how white matter integrity in ILF relates to functional activity during visual processing. Both Gschwind et al., (2012) and Pyles et al., (2013) report that white matter fibers connecting specialized regions of the face-processing network overlap with a portion of the ILF, suggesting that some fibers within ILF are important for transmitting neural activity related to face perception.

However, in the current study, no relationship was found between face-selectivity in fusiform face area and average FA of the entire ILF pathway, suggesting simply averaging the values for the entire tract—a common practice in DTI work (Bennett et al., 2012; Burzynska et al., 2010)—may not be the most sensitive predictor of brain activity within a small and localized region of the ventral visual pathway. Rather, FA within a more proximal region of ILF (mid-fusiform gyrus) was found to be the best predictor of face selectivity within FFA, suggesting that fibers within middle-fusiform portions of the ILF may subserve specialized neural activity in face-selective regions of cortex. Earlier behavioral work reports a selective association between inferior fronto-occipital fasciculus (IFOF; a fiber pathway passing through fusiform to frontal regions of the extended face-network) and better face discrimination, particularly for more difficult judgments, whereas ILF showed no such relationship (Thomas et al., 2008). Given that IFOF and ILF show substantial spatial overlap in fusiform regions (Pyles et al., 2013; Liu et al., 2020), it is possible that our mid-fusiform subsection contains fibers from both pathways, which might explain the specific relationship we find between mid-fusiform FA and FFA face selectivity. Future work is needed to examine a dissociation of mid-fusiform ILF and IFOF in predicting face selectivity in both core and extended regions of the face network.

On the other hand, whole ILF FA was found to be a significant predictor of selectivity for animacy across a large area of ventral visual cortex. Therefore, as a whole, white matter fibers of the ILF may subserve basic visual processing involved in superordinate category distinction within ventral visual pathway. Together, these results suggest that white matter exerts a local effect on neural activity, such that higher anisotropy in a small portion of ILF better predicts functional activity in proximal regions of cortex, whereas the entire ILF predicts broad differences in activity across ventral visual cortex. Although it is important to note that no standard exists for testing different gradients along white matter pathways using probabilistic tractography (see Yeh, Badre, & Verstynen, 2016), the current findings illustrate the utility in segmenting major white matter structures when corresponding functional regions are small and localized.

### Different properties of white matter microstructure predict functional selectivity

The current study used mean tract fractional anisotropy (FA) as the primary index of white matter microstructure to predict functional selectivity. FA is a scalar measure that describes the relationship between the rates of water diffusion parallel to the primary direction of diffusion (axial diffusivity) relative to diffusion in perpendicular directions (radial diffusivity). Therefore, while FA is a useful measure for describing overall shape and orientation of diffusion, it is also interesting to consider how changes in both primary (axial) and perpendicular (radial) diffusivity may be accounting for differences in FA. In general, studies examining effects of age on white matter find that age-related decreases in FA can be due to relative changes in both axial and radial diffusion. Specifically, decreased FA has been accounted for by three different patterns: (1) increased radial diffusivity only, which may indicate dysmyelination of the axonal sheath; (2) increased radial and axial diffusivity, which may indicate a more severe dysmyelination; and (3) increased radial and substantial decreases in axial diffusivity, which may indicate damage to the white matter axons (Bennett et al., 2010), although it is important to keep in mind that these are proxy metrics.

In the current study, decreased FA in middle-fusiform ILF predicted decreased selectivity for faces in FFA, and additional analyses found that decreased selectivity was also predicted by increased radial and decreased axial diffusivities (pattern #3), independent of the effect of age. This pattern of increased radial and decreased axial diffusivity may indicate that face selectivity in fusiform is sensitive to the pattern of diffusivity in nearby white matter which may be characterized by slight alterations to axonal cellular boundaries. These results suggest that white matter in middle-fusiform ILF may be more susceptible to more severe forms of degradation, and this in turn may impede functional response in nearby cortex. To note, the associations between white matter metrics and neural selectivity persisted beyond the effect of age, therefore differences in diffusivity likely represent subtle individual differences, rather than age-specific or age-dependent white matter degradation.

On the other hand, only increased radial diffusivity in whole ILF (pattern #1) was sensitive to decreased selectivity for animacy across ventral visual cortex, suggesting that dedifferentiation on a broader scale may be predicted by less severe differences in ILF white matter fibers. Furthermore, we report a significant age × ILF microstructure interaction (for both FA and radial diffusivity), characterized by stronger structure-function coupling in younger adults that weakened with age and was no longer significant after age 70. White matter in ILF matures through early adulthood, characterized by increased FA and decreased radial diffusivity, and peaks around ages 26-29 before beginning to decline (Westlye et al., 2010; Lebel et al., 2012). Furthermore, our sample indicates that ILF degradation becomes more severe after the age of 52 (**Figure 1**). Therefore, it is possible that this interaction is capturing the window with the greatest variability in individual differences in white matter maturity (for younger adults) and early and middle stages of white matter degradation (for middle-aged and older adults). That is, optimal white matter microstructure is most predictive of brain function at these transitionary points of the lifespan, but this relationship weakens as white matter aging becomes more prominent.

### Interpreting the current results in the context of prior DTI-fMRI aging studies

Prior studies examining the effect of age on the relationship between white matter microstructure and functional activity have largely examined how white matter is related to changes in magnitude or modulation of BOLD response during a cognitive task, with the largest focus on frontal regions of the brain (see Bennett & Rypma, 2013 for a review). Researchers generally report that decreased frontal white matter integrity in older adults corresponds with overactivation of frontal cortex (Burzynska et al., 2013; Daselaar et al., 2015; Davis et al., 2012; de Chastelaine et al., 2011; Hakun et al., 2015; Madden et al., 2007; Persson et al., 2006; but see also de Chastelaine et al., 2011 and Davis et al., 2012 for opposite findings), suggesting that the general age-phenomenon of increased activity in frontal cortex may be a compensatory response to impaired white matter degradation (Daselaar et al., 2015). White matter degradation in older adults has also been associated with failures to deactivate default regions during task (Brown et al., 2015) and failure to up-modulate prefrontal activity in response to increasing task demands (Webb et al., 2019).

We extend this prior work on higher-level cognition by moving our focus to structure-function relationships in ventral visual cortex during a basic perceptual task. Due to the highly organized and hierarchical nature of visual representations in ventral visual cortex (Grill-Spector & Weiner, 2014), we were able to target both focal (i.e., fusiform face area) and broad patterns of specialized functional activity. We report that dedifferentiation of functional response (i.e., less selective response) could be accounted for by decreased FA in underlying white matter at both fine-grain and coarse scales. Although this structure-function association can be interpreted as age-independent, examining the unique variance explained by age versus white matter microstructure in our regression models reveals that together these two predictors explained more variance in functional selectivity than on their own, suggesting both age and white matter play an important role. Furthermore, we find evidence for an age-dependent relationship between ILF FA and broad functional selectivity that is not significant in our oldest adults (ages 70+). This suggests that intact ILF microstructure is predictive of less dedifferentiated (i.e., more “youth-like”) functional patterns, but only to the point when white matter degradation is not too severe. Our findings, in conjunction with prior DTI-fMRI studies, may indicate that decreased white matter connectivity predicts maladaptive alterations in brain function, be it decreased selectivity, frontal over-recruitment, or failure to modulate functional activity. Therefore, underlying white matter structure is an important variable to consider when interpreting age-differences in brain function.

### Conclusions

In sum, the present study offers the first examination of how white matter microstructure (through multiple white matter indices) in the ILF predicts functional selectivity in the ventral visual cortex across the entire adult lifespan. This study provides a comprehensive view of the relationship between measures of white matter microstructure and selectivity in a healthy, aging population, which may help dissociate normal from pathological brain aging in future studies. There are several important points to take away from these findings. First, the effect of white matter on functional selectivity appears to be local, such that neural activity is best predicted by integrity in nearby white matter. Specifically, whole ILF predicts selectivity for animacy across the entire ventral visual cortex, whereas a smaller, proximal region of ILF predicts selectivity for faces in a focal region of fusiform gyrus. Next, coupling between ILF white matter and broad functional selectivity for animacy is not steady across the lifespan—rather, white matter is the strongest predictor of neural selectivity at younger ages, and this relationship weakens, but is maintained until age 70. Future longitudinal studies directly examining intra-individual changes in white matter tracts in relation to functional brain activation are needed to confirm the cross-sectional findings reported here.

## Acknowledgments

This study was supported in part by National Institutes of Health grants: R37-AG-006265 (DCP); R37-AG-006265-S1 (DCP); R01-AG-056535 (KMK); R01-AG-057537 (KMR).

## Footnotes

1 To note, mid-fusiform FA remained predictive of FFA selectivity after also accounting for FA in two other segments of ILF more distal from FFA (one approximately 15mm posterior, and one approximately 15mm anterior; *F*(1,272) = 4.67, *p* = .032), suggesting that these effects are specific to mid-fusiform ILF due to its proximity to the functional region of interest, rather than due to inherent regional differences in FA values along the ILF.

